# Simulations of *CYP51A* from *Aspergillus fumigatus* in a model bilayer provide insights into triazole drug resistance

**DOI:** 10.1101/088468

**Authors:** Anthony Nash, Johanna Rhodes

## Abstract

Azole antifungal drugs target *CYP51A* in *Aspergillus fumigatus* by binding with the active site of the protein, blocking ergosterol biosynthesis. Resistance to azole anti-fungal drugs is now common, with a leucine to histidine amino acid substitution at position 98 the most frequent, conferring resistance to itraconazole. In this study, we create a homology model of *CYP51A* using a recently published crystal structure of the paralog protein *CYP51B*. The derived structures, wild-type and L98H mutant, are positioned within a lipid membrane bilayer and subjected to molecular dynamics simulations in order improve the accuracy of both models. The structural analysis from our simulations suggests a decrease in active site surface from the formation of hydrogen bonds between the histidine substitution and neighbouring polar side chains, potentially preventing the binding of azole drugs. This study yields a biologically relevant structure and set dynamics of the *A. fumigatus* Lanosterol 14 alpha-demethylase enzyme and provides further insight into azole antifungal drug resistance.

## Introduction

*Aspergillus fumigatus* is the causative agent of invasive aspergillosis (IA), a life-threatening infection in immunocompromised individuals. Orally administered azole antifungals are the first-line drugs for the treatment of IA in humans, however, resistance to these drugs is rising. Azole-resistant *A. fumigatus* strains have been identified across the globe [1, 2, 3, 4], with resistance to the drugs itraconazole common, and resistance to voriconazole increasing, presenting an evolving problem for public health [5, 6].

Mutations in the target gene of azole anti-fungal drugs, *cyp51a*, have been shown to confer resistance phenotypes in *A. fumigatus* [7, 8, 9]. *cyp51a* encodes a CYP450-dependent enzyme, 14*α*-lanosterol demethylase, which is inhibited when azole drugs bind to the catalytic iron of the heme prosthetic group in the *CYP51A* active site. An amino acid change from leucine to histidine at position 98 of *cyp51a* is the most common resistance mechanism [10], conferring resistance to the first-line antifungal drug itraconazole.

Structural and biochemical characterisation of fungal *CYP51* is difficult due to being an embedded membrane protein. Previous studies have used crystal structures of *Mycobacterium tuberculosis CYP51* [11] (*MT-CYP51*)[12, 13] and non-fungal eukaryotic species [14], with low sequence similarity to fungal *CYP51* [15]. Bacterial *CYP51* proteins are also soluble, whereas fungal *CYP51* is instead an integrated membrane protein.

Snelders *et al.* [16] improved the homology model by using the crystal structure of the human lanosterol 14*α*-demethylase (PDB code: 3I3K), conferring 38 % sequence identity. Although Snelders *et al.* acknowledge the presence of the heme prosthetic cofactor in *CYP51A*, there is no description of heme in their molecular dynamic (MD) simulations. Therefore, the results presented in Alcazar-Fuoli *et al.* and Snelders *et al.* do not represent the true extent of the biological problem.

The *A. fumigatus* protein *CYP51B* (*AfCYP51B*) has 59% amino similarity with its paralog, *CYP51A* (*AfCYP51A* ), with the X-ray crystal structure for *AfCYP51B* published in 2015 [17]. It is thought that *AfCYP51B* is expressed constitutively, whilst *AfCYP51A* expression is induced by the presence of azole drugs; however, no redundancy in function has been observed when *Afcyp51a* has been knocked-out. A recent homology study by Lie *et al.,* [14], yielded stability measurements of three azole drugs interacting within an implicit solvated buried active site of *AfCYP51A*. A set of model homologs that demonstrated superior coverage quality were subjected to an energy minimisation step. Minimised structures, more often than not, represent an energy local minimum. The sufficient exploration of phase-space in pursuit of a more biological comparable structure requires extensive time-integration of velocities assigned to the structure. A crystallographic study on cytochrome P450 suggests constraints that orient the catalytic domain relative to a bilayer [18]. The previous computational studies had excluded the complex micro-environment of *AfCYP51A*, notably, the N-terminal anchored across a lipid membrane bilayer.

We employ an all-atom representation of *AfCYP51A* using the crystal structure of *AfCYP51B* as the homology model template. From this structure, a L98H single point mutant variant can be created. The transmembrane (TM) domain was identified and the integral TM domain *α*-helix was embedded in a model 1-palmitoyl-2-oleoyl-sn-glycero-3-phosphocholine (POPC) membrane bilayer. The inclusion of a Fe^2+^ dummy model provides the simulation with the dynamic exchange between the metal and its immediate environment and is a significant improvement over earlier models of covalently-bound metal models [19]. The cationic dummy atom approach describes a metal centre coordinated by a set of cationic dummy atoms. This approach not only captures correct electrostatic effects but it is also capable of dynamic ligand exchange with the environment and maintaining the correct number of ligand donors [20]. MD simulations performed in this study further the biological relevance and understanding of azole antifungal drug resistance from the single amino acid mutation, L98H, far from the buried active site [21, 22].

## Computational methods

### Model generation

All raw reads and relevant information in this study have been submitted to the European Nucleotide Archive under project accession number PR-JEB8623. Genomic DNA sequences for both the L98H mutant and wild-type *Afcyp51a* were obtained from a previous study [7]. A three-dimensional homology model, one for the wild-type and another for the L98H mutant, was produced within Schrodinger Maestro (2015-4) using a BLAST homology search. The inclusion threshold was set to 0.005 and three iterations were applied. The first thirty-six residues from the N-terminal and last five residues before the C-terminal were not predicted by the BLAST homology search. The derived tertiary structure from the BLAST homology search also included the heme prosthetic group and Fe^2+^ metal ligand, coordinated to a proximal cysteine thiolate.

### Simulation details

All-atom MD simulations were performed using the GROMACS simulation package, version 5.1.2 [23]. The integration time step was set to 2 fs, and the leapfrog algorithm was used for time-step integration. All bonded atoms were holonomically constrained using the LINCS constraint algorithm [24]; the LINCS iteration and order was set to 1 and 12, respectively. The short-range neighbour interaction list cut-off was fixed to 1.2 nm and updated ever 10 fs. Short-range interactions were modelled using a 12-6 Lennard-Jones potential and were truncated at 1.2 nm using the potential-shift cutoff modifier from 1.0 nm. Short-range electrostatic interactions were truncated at 1.2 nm and long-range electrostatics were calculated using the Particle Mesh Ewald scheme [25] with dispersion correction applied to the energy and the pressure.

The temperature was coupled using the V-rescale scheme [26] over two groups; the protein-complex, including the protein, heme prosthetic group and Fe^2+^ metal-ligand; and the solvent with counter ions, at a physiological value of 310 K. The pressure was maintained at one atmosphere by first applying a semi isotropic Berendsen coupling [27] before switching to the Parrinello-Rahman scheme [28], to increase the accuracy of an NPT (fixed number of particles, fixed pressure and fixed temperature) ensemble. Loose coupling was applied between the protein-complex and the water environment and the compressibility was set to 4.5e-5. Periodic boundary conditions were applied in three dimensions. Coordinates, velocities and energies were saved every 5 ps during production simulations.

A cubic periodic unit cell was employed and the solute was fully hydrated with Tip3p water solvent molecules. The distance of the solute to the unit cell boundary was adjusted to prevent periodic image artefacts. Calcium ions were added to yield a net neutral charge.

### Model POPC bilayer construction

A united-atom POPC lipid representation was employed. POPC lipids are naturally present in eukaryotic cell membranes and parameters provided [29] show reliable free energy estimates compatible with the Amber99SB force field [30]. A three-dimensional structural representation of a single POPC lipid was replicated in the x-y plane to make a slab of 126 lipids. The structure was copied, flipped and displaced to align lipid tail ends, resulting in a 252 lipid bilayer. The system was fully solvated with Tip3P water molecules. Using the simulation details provided above, the structure was subjected to a steepest descent minimisation with a maximum force tolerance of 1,000 kJ mol^−1^ nm^−1^. Then, a 1,000 kJ mol^−1^ position restraint was applied to the phosphate group of each lipid before subjecting the system to a 100 ps simulation using the NVT ensemble. The position restraint was removed, and a semi isotropic Berendsen pressure coupling was added for a 1 ns simulation. Finally, the Berendsen coupling was replaced with the Parrinello-Rahman scheme and the system was left to equilibrate for 30 ns. Measurements of the average bilayer thickness and area per lipid indicated that the final structure was suitable for protein insertion.

### Protein structural equilibration

The protein-complex was energy minimised in vacuum using steepest descent with a maximum force tolerance of 1,000 kJ mol^−1^ nm^−1^. The resolved structure was then fully hydrated with solvent molecules and necessary counter ions, before subjecting the complete system to a further energy minimisation whilst applying position restraints on the protein-complex. The restraints were then removed from the protein, and the structure was minimised further. Finally, all restraints were removed and the system underwent a final steepest descent minimisation.

The atomic-partial charges, bond force constants and bond equilibrium values for the heme prosthetic group were taken from [31]. The non-bonded potentials, force constants, equilibrium values, and atomic-partial of the Fe^2+^ dummy atom were taken from [20]. The non-bonded potentials were fitted to the Gromacs sigma and epsilon format using the method outlined in the supporting information of Liao *et al.* [32]. Observation of Fe^2+^ bond distances revealed bond distances beyond the equilibrium bond value. An MD simulation using the NVT ensemble, with the temperature set to 274 K and an integration step to the smaller value of 0.001 fs, was performed for 10 ps to rectify this discrepancy. Position restraints of 1,000 kJ mol^−1^ were then applied to both carbon atoms immediately bonded to the heme nitrogen atoms along with the central Fe^2+^ atom; the dummy atoms were left unrestrained. The integration step was set to 0.002 fs, from hereafter, and a 50 ps NPT simulation using the Berendsen pressure-coupling was performed. The coordinates and velocities were preserved and the simulation was extended by a further 50 ps using the Parrinello-Rahman pressure coupling scheme.

### Protein insertion

The identified TM domain was positioned across the hydrophobic region of the lipid bilayer. The intra-celluar N-terminal sequence was positioned within the phosphate head group and lipids that overlapped protein residues were removed. A set of 1,000 kJ mol^−1^ position restraints was then applied to the protein, heme prosthetic group and dummy iron and the system was subjected to a steepest descent minimisation. The restraints were maintained and a 50 ps simulation using the NVT ensemble was performed to initiate the packing of lipids around the protein. The bilayer-protein complex was then fully solvated using Tip3P molecules and neutralised using counter-ions. Water molecules that had been automatically placed within the lipid alkyl tails were removed.

## Results and discussion

### *AfCYP51A* homology modelling

A BLAST sequence search was performed using globally conserved residues matching the Cytochrome P450 family revealed 221 homologs, from which the top-scoring crystal structure, sterol 14-*α* demethylase (*AfCYP51B*), from the pathogenic filamentous fungus *A. fumigatus* in complex with the small molecule (R)-N-(1-(2,4-dichlorophenyl)-2-(1H-imidazol-1-yl)ethyl)-4-(5-phenyl-1,3,4-oxadi-azol-2-yl)benzamide [17] (4UYL), yielded an identity of 65 %. Alignment was performed using Clustal W [33], and the secondary structure and loop structures was calculated using Prime [34]. The heme prosthetic group and Fe^2+^ ligand were included. An earlier study by Liu *et al.* [14] had identified the top ten homologs, of which the maximum identity scored 50.7 %. Since that study, the crystal structures 4UYL and 4UYM of *CYP51B* from *A. fumigatus* have been made available, resulting in the additional sequence identity captured in this study.

The homology search yielded an N‐ and C-terminal truncated three-dimensional structure. To reconstruct the two missing sequences, an I-TASSER calculation was performed over both sequences. This is a free protein-fold predictor service that has been shown to be consistently grouped among the most accurate services available [35]. The prediction of the wild-type and L98H mutant protein structure yielded identical confidence and TM-scores, 1.44 and 0.91±0.06, respectively. The confidence intervals spans [-5,2], and the higher the value the more confidence, whilst a TM-score >0.5 indicates a model of correct topology. When superimposed over the homology model, the I-TASSER prediction had produced an identical structure but with the addition of the missing terminal segments. Given that I-TASSER does not transfer co-factors, prosthetic groups or metal ions, the BLAST homology model was used and the terminal sequences from the I-TASSER results were concatenated. A clear similarity between our complete homology model over the top scoring 4UYL template can be seen in Figure 1.

**Figure 1:**
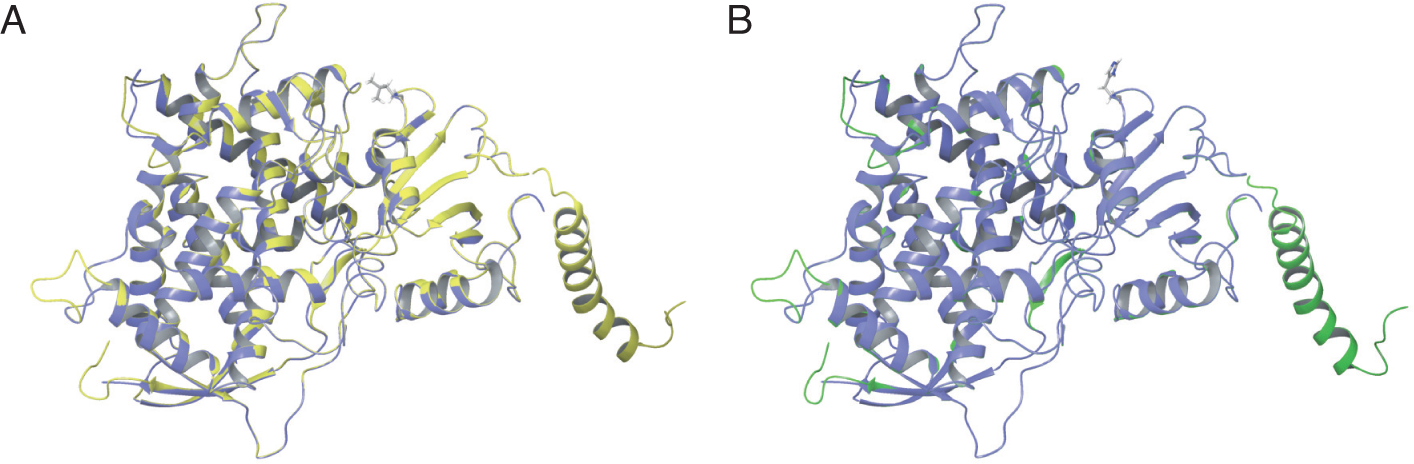
The homology models including N‐ and C-terminus concatenated using I-TASSER structural predictions. Both wild-type (A) and L98H mutant (B), in yellow and green, respectively, have been superimposed over the *AfCYP51B* template structure, blue.

### Membrane bilayer molecular equilibration

The accurate representation of the interactions between a lipid bilayer and an integral membrane protein, whether as an anchor to a bilayer or as an extensive membrane channel, is only possible within the context of explicit lipid representation. Established studies using implicit bilayers were able to account for the hydrophobic mismatch between TM domains and the system solvent [36], however, due to membrane deformation from lipid-protein rearrangement, the tight-coupling between protein structure and protein function can only be represented by explicit lipid bodies [37]. The grand average of hydropathy (GRAVY) [38] value for the wild-type and L98H mutant were -0.216 and -0.230, respectively. A comparison between the absolute terms do not reveal a great deal of hydropathy difference between sequences, However, as a pair, they are considered hydrophobic. This is indicative of their role as integral membrane proteins[18], a characteristic not seen before in AfCYP51 computational chemistry studies.

Crystal structures of *Saccharomyces cerevisiae* CYP51 have identified an *α*-helical integral membrane anchor segment leading from the N-terminus [18]. Using TMHMM [39], we were able to predict the spatial arrangement between the bilayer and the protein (SI Figure 2). The first six residues at the N-terminal region of wild-type and L98H mutant were predicted to be a random coil and exposed to the intra-cellular region of the bilayer. This was immediately followed by a 23 residue long *α*-helical integral TM domain. Two coils, similar in length, are expected to interface with the lipid chains, separated by very short extra-cellular sequences, M39 ‐ F61 and I71 ‐ E91, as the sequence is brought back into the bulk of the protein. The relationship between our *AfCYP51A* all-atom models with the lipid bilayer differs in luminal bound peptide segment length with that of the *Saccharomyces cerevisiae CYP51* crystal structure [18]. However, at closer examination between species, the *CYP51* differs in primary sequence length. Further, the TM domain predictions over the *AfCYP51A* sequence did not yield a TM domain delimited by an intracellular followed by an extracellular region. In addition, during the production MD simulations of wild-type and L98H mutant, the suggested TM domain remained helical and continued to span the bilayer. Finally, the anchoring of the N-terminal region across a lipid bilayer prevents those residues from uncoiling in the polar solvent environment (or potentially under vacuum) and interacting with regions of the catalytic domain, making for a very inaccurate structural representation. Our earlier simulations of *AfCYP51A* presented solely in water solvent (not presented) demonstrated the immediate uncoiling of the TM domain.

We explored the validity of the predicted TM domain further by subjecting the sequence to an experimentally derived free energy of membrane insertion calculation, ΔG predictor [40]. In short, the predicted ΔG of amino acid insertion into the Endoplasmic Reticulum membrane by means of the Sec61 translocon was used as a measure of identifying suitable TM domain sequences. The accumulation of individual amino acid contributions (see SI Figure 3) yielded -0.50 kcal/mol. The non-polar side chains of leucine, isoleucine, valine, and phenylalanine contribute favourably to membrane insertion, whilst alanine, despite being a non-polar residue, incurs a slight penalty. Asparagine incurs the greatest free energy penalty, however, statistically, asparagine is often found within the centre of a TM domain *α*-helix [40]. Taking into account the contribution from the hydrophobic moment, +0.56 kcal/mol, and the contribution for the helix length, -1.20 kcal/mol, the total propensity for insertion was a favourable -1.14 kcal/mol.

Once the TM domain was spanning the POPC lipid bilayer and both systems had been fully solvated (see methods), a series of equilibrium steps were performed. The heme prosthetic group and metal iron were restrained as described, and 1000 kJ/mol position restrains were applied over the protein heavy backbone atoms. The system was subjected to a 10 ps simulation using the NVT ensemble. The restraints were maintained and the Berendsen pressure-coupling was applied to a simulation of 100 ps. Reducing the restraints to 100 kJ/mol the pressure-coupling was replaced with the Nose-Hoover coupling and a further 100 ps was applied. The system continued for a further 100 ps but with a reduced position constraint of 10 kJ/mol.

NPT production runs of 65 ns and 45 ns, were performed for the wild-type and L98H mutant systems, respectively, from which the root mean square deviation (RMSD) and the area per lipid (APL) were calculated. The RMSD of the initial 40 ns, and 20 ns of the wild-type and L98H mutant, respectively, suggest that both systems were still undergoing structural adjustment (SI Figure 7). It is well understood that computational bilayer systems are prone to long velocity auto-correlation times and therefore require significant equilibration times [41]. Visual observation of both systems reveals a stable tertiary structure, with the TM domain consistently spanning the bilayer (Figure 2 A) and the N-terminal exposed to the solvent (Figure 2 B). The secondary structure of the TM domain remained stable (Figure 2 C) and the channel to the buried active site remained accessible to the solvent environment (Figure 2 D).

**Figure 2:**
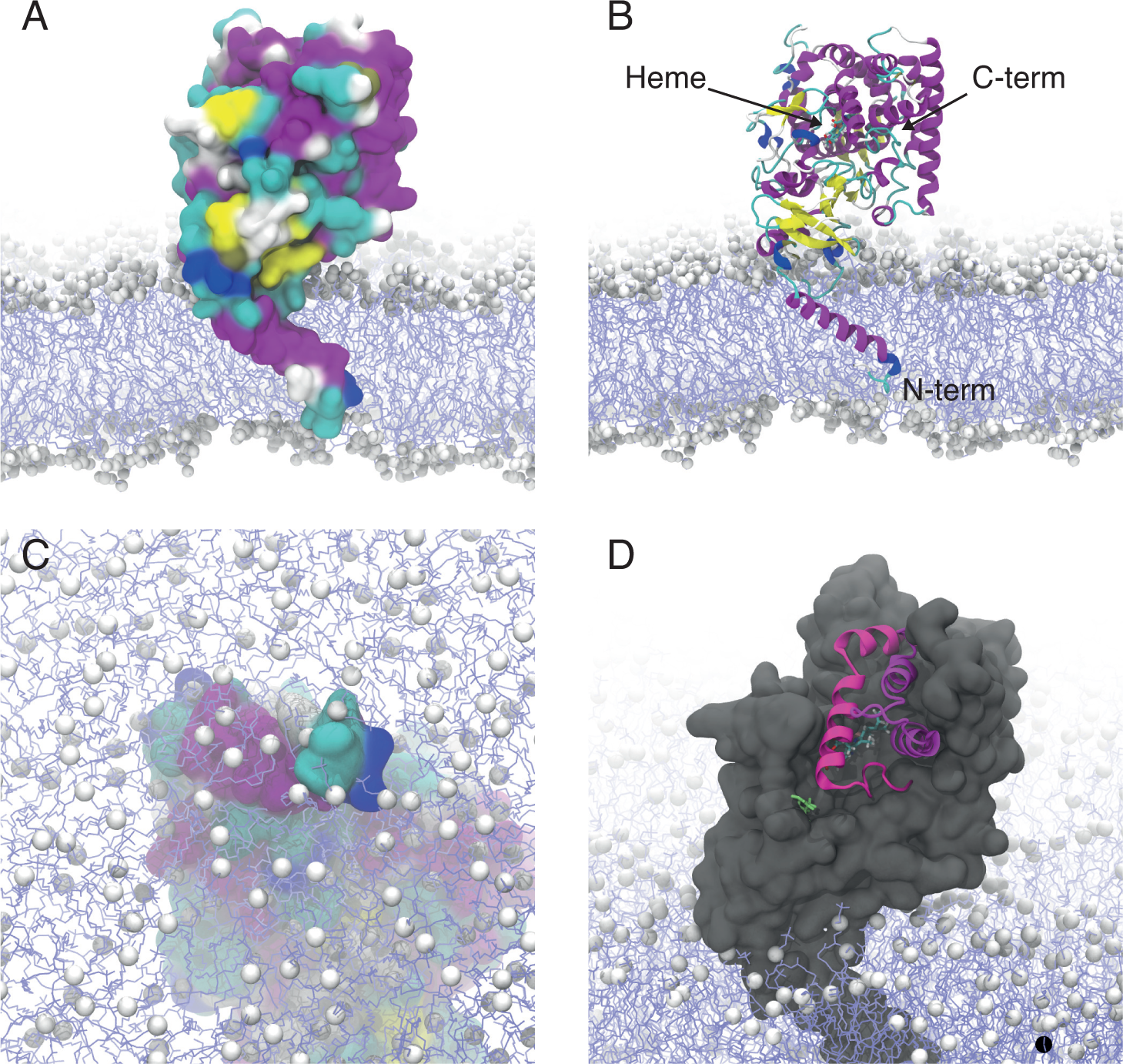
Snapshot representative models (taken from the final frame, 45 ns, of the *AfCYP51A* L98H mutant) of *AfCYP51A* anchored to a model POPC lipid bilayer. White space-filler spheres represent lipid head group heavy atoms and light blue lines represent alkyl chains. Water molecules have been removed for clarity. (A) The bulk of the protein using van der Waals distances. (B) A depiction of the protein secondary structure. (C) The N-terminus as seen passing through the lipid head group region. (D) The entrances to the active sight (pink helices) nearest to the L98H mutation site (lime green). The heme prosthetic group with bound Fe^2+^ can be distinguished from the bulk of the protein.

Adjustment to the lipid bilayer can be monitored by comparing the APL to experimental values. Frames for lipid analysis from both systems were retained every 5 ns (SI Figure 4 and SI Figure 5). After the short initial equilibrium steps, the average APL in both membranes was approximately 76.5 Å^2^, well beyond the experimentally derived value of 68.3 Å^2^ for POPC [42]. Within the first 5 ns, the APL of both membrane bilayers drops to approximately 67 Å^2^, and after which the value continually fluctuates within good agreement with the experimental value. In the presence of the TM domain, both sets of boundary lipids (those lipids that immediately interface the protein) show a decrease in APL, indicative of an ordered configuration to the hydrocarbon chains about the *α*-helical TM domain. The morphology in hydrophobic thickness of the lipid chains is a thoroughly studied consequence of the hydrophobic mismatch between an integral membrane protein and its membrane bilayer [43].

### *AfCYP51A* molecular dynamics simulation

Structural analysis of the L98H mutant and a comparison to the wild-type was performed using the last 25 ns of each respective production run, from which, the coordinates and velocities of each system at 5 ns intervals were obtained for visual inspection. The comparison in Ramachandran plots between the final frame (65 ns) of the wild-type sequence with the respective homology model initial structure (SI Figure 1) yields a significant alterations to the dihedral angles of the protein backbone. This is an initial indication that a homology model falls short of reporting an accurate representation of *AfCYP51A*.

To the H98 substitution and native L98, functional groups of side chains R451, N125, E104 and the carbonyl group on the backbone of C123, were seen as potential sites for protein-fold interactions. The aliphatic side chain of L98 and functional ring of HIS98 were used as a centre-of-mass reference for a distance calculation between the aforementioned groups over 25 ns of both trajectories. Non-bonded inter-atomic interactions, considering the offset from a centre-of-mass calculation, can be seen within 0.6 Å(SI Figure 8). During the 25 ns of the wild-type simulation, N125 and C123 were consistently beyond inter-atomic bonding. This came of little surprise given the hydrophobic mismatch between the functional group of cysteine and asparagine with the aliphatic side chain of leucine. Using the saved velocities and coordinates from the five retrieved time frames (40 ns to 65 ns at 5 ns intervals), 40 ps continuation simulations were recalculated whilst preserving the trajectory and coordinate frames were captured every 0.2 ps. Time lapse representations over the duration of the short simulations starting from each 5 ns interval time frame have been drawn in Figure 3. The native L98 is consistently beyond the interaction of the N125 carboxamide functional group, which in turn, is shown to interact with the polar complex guanidinium of R451. The native L98 side chain can be observed consistently orientated into the aliphatic region of the side chain of N125. The distance between L98 and E104, although not too great to contribute towards an interatomic interaction would be under a lot of repulsion from the negative charge associated with the *α*-carboxylic acid group of glutamic acid. A centre-of-mass distanced based measurement was performed between the substitution H98 and the functional groups of the nearest side chains R451, N125, E104 and the carbonyl group on the backbone of C123 (SI Figure 8). As with the wild-type, each 5 ns interval performed a 40 ps rerun of the trajectory, whilst recording frames every 0.2 ps (Figure 4). The polar complex guanidinium of R451 was shown to sit far within the interaction range of the partially protonated imidazole side chain of histidine, however, given the distance-offset associate with the centre-of-mass geometry, it is possibly that there was a brief association towards the latter 10 ns. The functional group of N125 was consistently within the short interaction distance with H98, establishing a hydrogen bond between the imidazole nitrogen bound hydrogen atom with the carbonyl oxygen of the asparagine side chain functional group. The interaction of the L98H mutant with the *α*-carboxylic acid group of glutamic acid and the carbonyl group on the backbone of the cysteine residues, are brief and present whilst the distance with N125 increases marginally. Unlike in the case of the wild-type, the L98H mutant can contribute to the stability of the neighbouring structure by forming hydrogen-bond interactions with water molecules, instigating bridging interactions between water molecules and close polar side chains.

**Figure 3:**
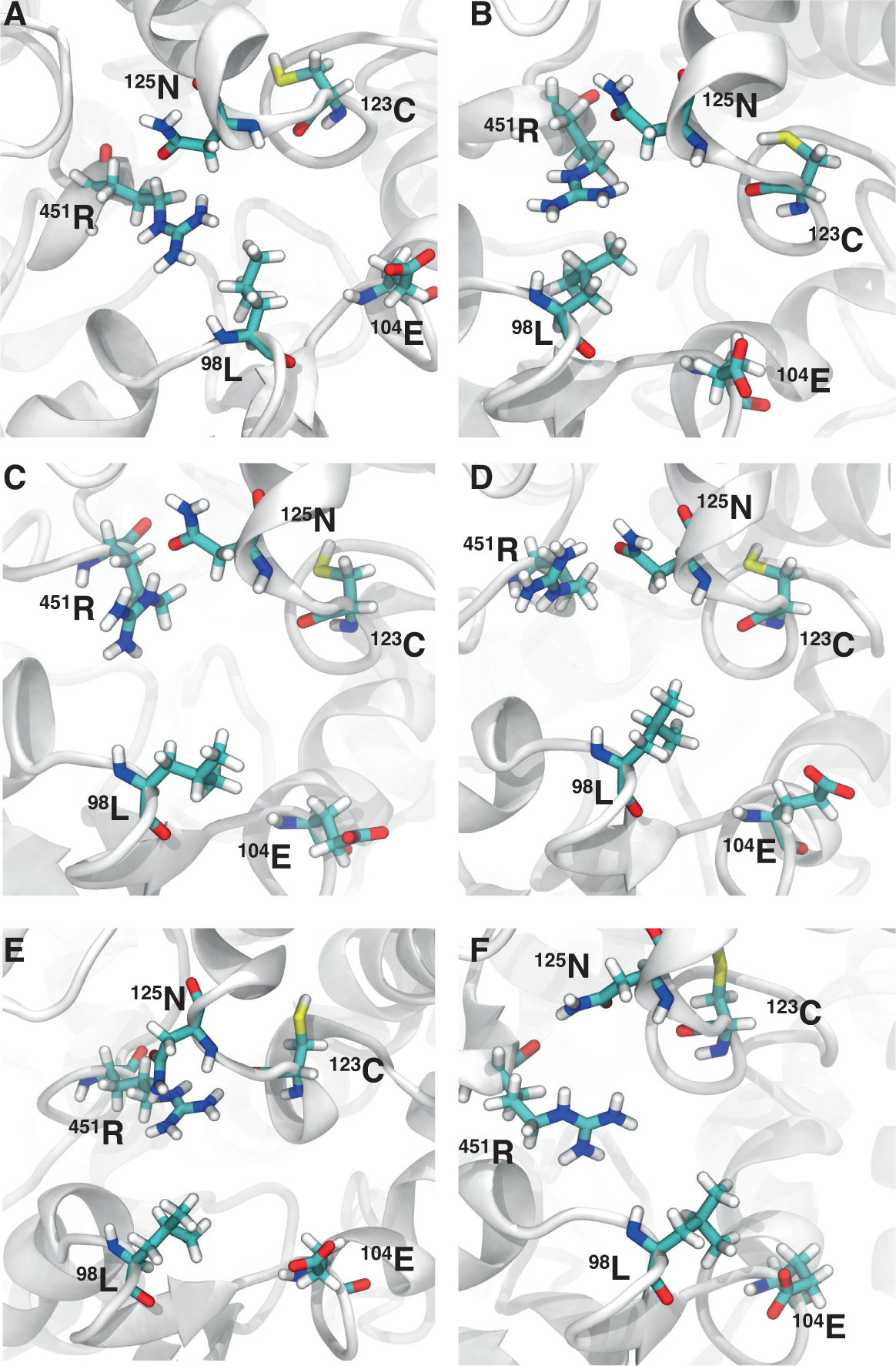
The structural arrangement of the wild-type L98 side chain with neighbouring side chains from a centroid frame calculated over the 40 ps simulations. Images A to F correspond to representative structural frames taken over 5 ns intervals from 40 ns to 65 ns. The secondary structure is represented in white ribbons and side chains in thick-coloured atom types.

**Figure 4:**
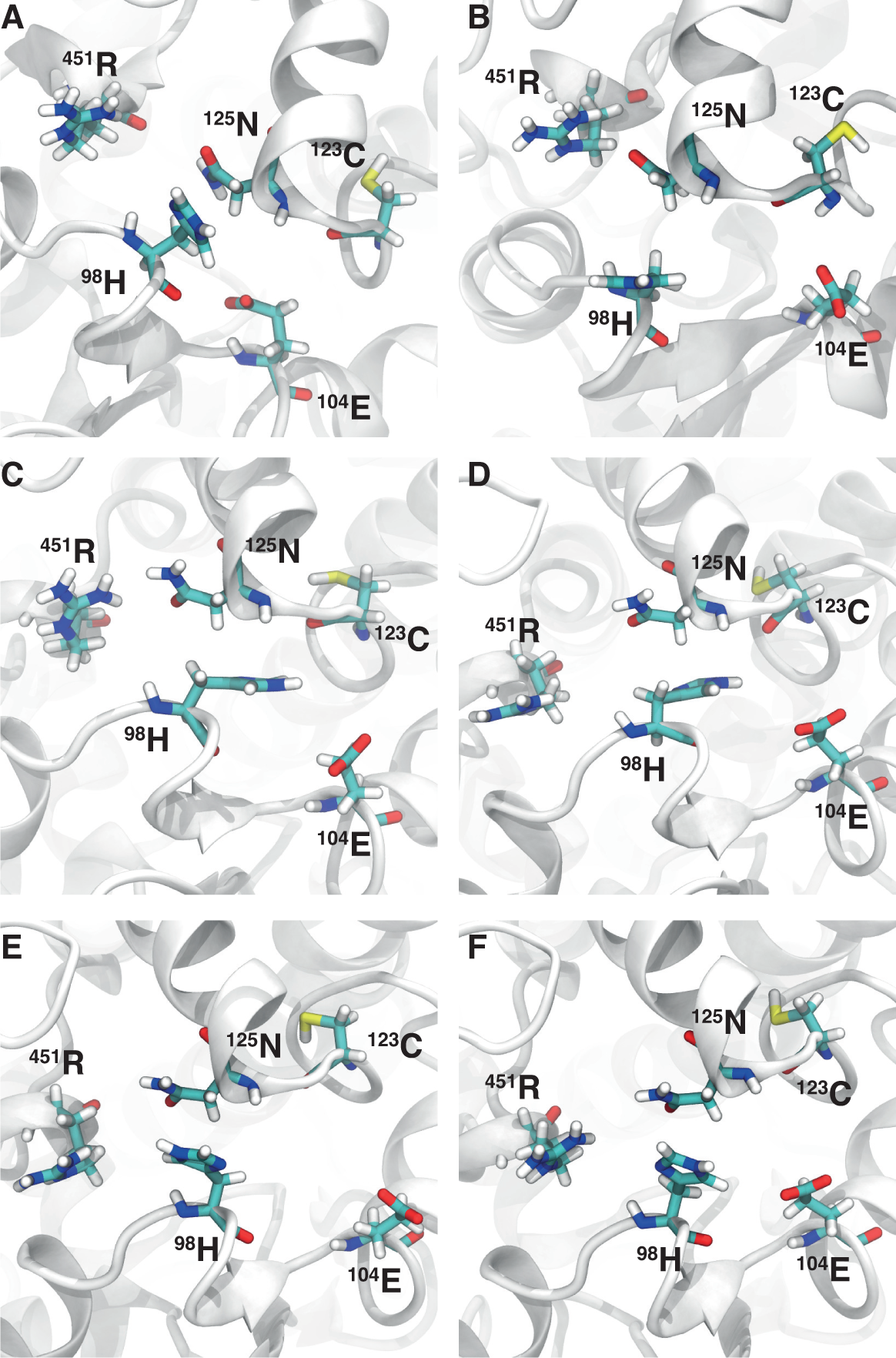
The structural arrangement of the mutant H98 side chain with neighbouring side chains from a centroid frame calculated over the 40 ps simulations. Images A to F correspond to representative structural frames taken over 5 ns intervals from 20 ns to 45 ns. The secondary structure is represented in white ribbons and side chains in thick-coloured atom types.

### Active site dynamics

The L98H substitution, far from the active site, clearly demonstrates a modification to the structural configuration of the neighbouring residues. From a visual inspection of the secondary structure arrangement, a change in side chain hydrophobicity, from non-polar to polar-charged, resulted in a contraction of the protein. The surface areas occupied on the outside of the catalytic domain of the residues H98, R451, N125, C123, and E104 is reduced as a result of a closer association between the substitution and the neighbouring polar side chains. This suggests, a looser structural configuration in the wild-type compared with the mutant, supporting the idea that azole-drug binding is reduced bu the mutation. The *α*-helix, denoted H1 in Figure 5, shifts closer to the channel entrance coil segment, C1, contracting the shorter H3 *α*-helix. The association between H98 and N125 leads to a short uncoiling of H2 *α*-helix nearest to the C2 coil.

**Figure 5:**
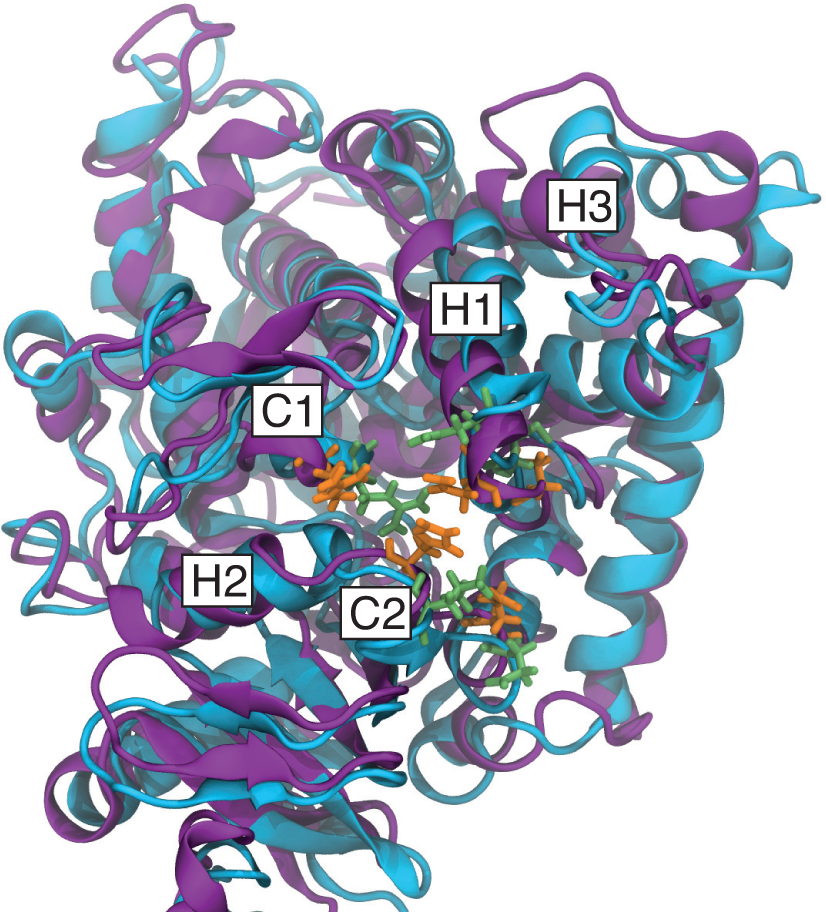
A secondary structure representation of *AfCYP51A* from the final frame of the wild-type (blue), and L98H mutant (purple). The *α*-helices and coils immediately adjacent to the single point mutation have been labeled. The side chains identified from interatomic coordination with the L98H single point mutant have been highlighted in orange, alongside their wild-type positions in green.

The adjustments to the secondary structure were considered in relation to the accessible active site surface of the wild-type and L98H mutant sequence. The surface area of the active site was calculated for any residue-ligand within 5 Åof the heme prosthetic group and coupled ion. Measurements were taken at 5 ns interval over the final 25 ns of each respective production run. The measurements, collected in Table 1 demonstrates a noticeable decrease in active site surface of the L98H mutant across all recorded instances compared to the wild-type.

**Table 1:** The active site surface area (Å^3^) of the wild-type and L98H mutant. Measurements were taken at 5 ns intervals across both final 25 ns trajectories.

| Binding site surface ($\text{\AA}^3$) | | | | | | |
| --- | --- | --- | --- | --- | --- | --- |
| Wild-type *AfCYP51A* | 40 ns | 45 ns | 50 ns | 55 ns | 60 ns | 65 ns |
|  | 1383.6 | 1221.5 | 1305.5 | 1298.3 | 1395.5 | 1430.7 |
| L98H Mutant *AfCYP51A* | 20 ns | 25 ns | 30 ns | 35 ns | 40 ns | 45 ns |
|  | 1075.9 | 1057.4 | 1020.2 | 1087.8 | 1122.6 | 1149.6 |

Ligand interaction diagrams were generated for each retrieved interval using Maestro 2015-4, highlighting the coordination of Fe^2+^ with the four pyrrole nitrogens within the heme plane, and the coordination of protein ligand donors to the polar groups of the heme prosthetic group. The contraction of the protein secondary structure from the substitution L98H resulted in a modification to the heme-protein interaction. In the wild-type (Figure 6), Y121 established a hydrogen bond coordination between the functional hydroxyl group of the tyrosine functional group with one of two of the carbonyl groups on the heme prosthetic group. Equally, Y107 established a similar coordination over half of the monitored production run, and the back bone amine hydrogen remained attracted to the negative charge of a heme prosthetic group oxygen. The guanidinium functional group of R369 was shown to coordinate with a set of hydrogen bonds to the oxygen atoms of the heme prosthetic group, during the first 10 ns. This coordination is lost after the final 15 ns. The final coordination is seen between the positively charged functional group of K132 with an oxygen atom of the heme prosthetic group, throughout the recorded wild-type production run. These recorded heme-ligand side chain interactions compare well to those observed in the *AfCYP51B* template crystal structure, notably Y122, Y126, R378, and K147 [17] (the difference in residue number is due to the offset caused by a difference in primary sequence length).

**Figure 6:**
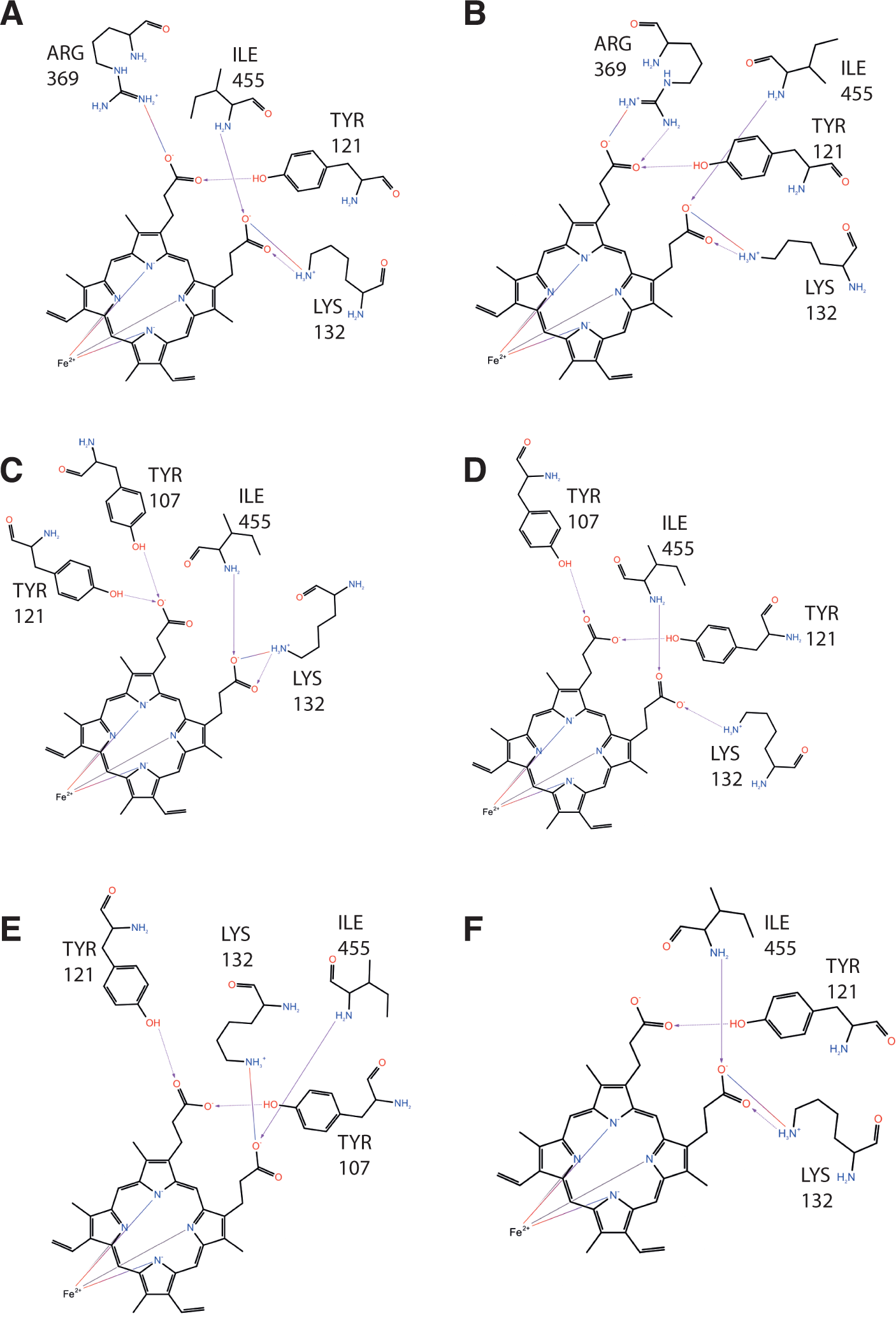
Ligand interaction diagrams for the heme binding site of the wild-type structure at each 5 ns interval (from A to F) over the final 25 ns trajectory. The binding pocket is indicated by a line drawn around the heme. Residues ligands are labeled explicitly and shown in stick-form. Non-polar in green, positively charged polar in purple, non-polar neutral (glycine) in yellow, neutral polar in light blue, an3d5 negatively charged polar in red

The reduced active site surface as a result of the L98H substitution, yield a set of heme-ligand interaction diagrams (Figure 7) different to those of the wild-type. Neither tyrosines or the arginine, seen during the wild-type coordination were present, and in their place, G456 was shown to form a hydrogen bond between its amine backbone with a carbonyl oxygen atom of the heme prosthetic group.

**Figure 7:**
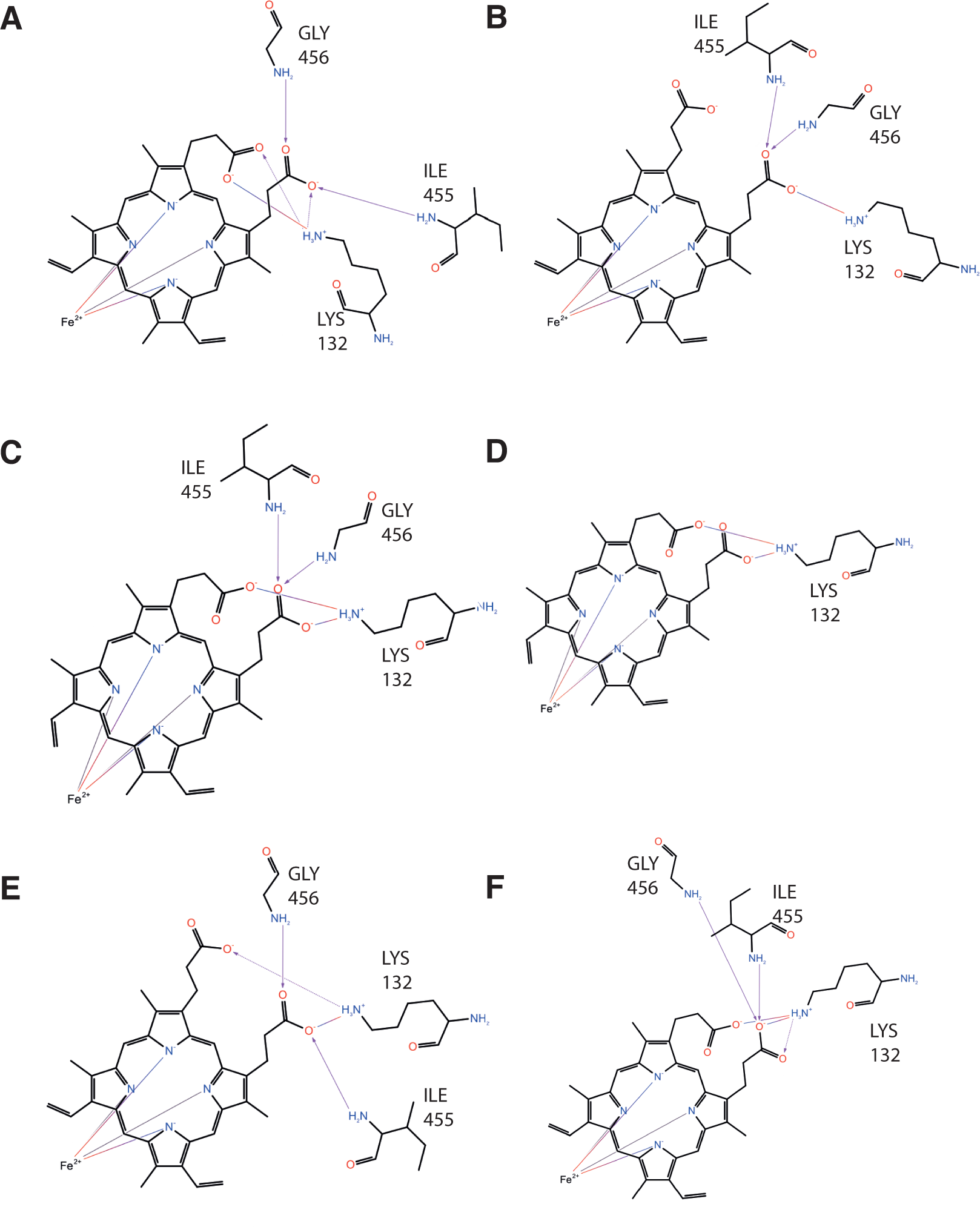
Ligand interaction diagrams for the heme binding site of the L98H mutant structure at each 5 ns interval (from A to F) over the final 25 ns trajectory. The binding pocket is indicated by a line drawn around the heme. Residues ligands are labeled explicitly and shown in stick-form. Non-polar in green, positively charged polar in purple, non-polar neutral (glycine) in yellow, neutral polar in light blue, and negatively charged polar in red.

Not only has the L98H amino acid substitution resulted in a contraction to the secondary structure leading to a reduced active site surface, but the protein coordination around the heme prosthetic group has changed significantly. It is within reason to suggest that these conformation changes may affect the nucleophilic nitrogen of the azole heterocyclic from acting as the sixth coordinating ligand with the heme ferric iron [44]. Furthermore, the structural changes may well affect the initial passage and subsequent binding of azole drugs into the buried active site.

## Conclusion

A similar study in 2011, by Snelders *et al.* [16], demonstrated a structural contraction to the active binding cavity and a deviation in ligand donors to the heme prosthetic group. Interestingly, the heme-ligands identified were different to those present in this study. Further more, the earlier study used a human template with a sequence identity of only 38 %. Our study has given further biological relevance with the recent availability of the *AfCYP51B* crystallographic structure, a paralog to our target. The model includes a TM domain that we integrated across a POPC lipid membrane bilayer, and with the inclusion of a dummy metal ion bound to a heme prosthetic group dynamic exchange between the Fe^2+^ and the immediate environment was possible.

Compared with earlier homology models [14], the inclusion of the integral membrane *α*-helix spanning an explicit lipid bilayer, enables time integration of the structure whilst circumventing the N-terminal region from uncoiling and interacting with the catalytic domain. Our efforts have yielded a very accurate and biologically relevant model of the *A. fumigatus* Lanosterol 14 alpha-demethylase enzyme.

Molecular dynamics simulations of the wild-type and L98H mutant revealed a measurable variation of interaction distance between the mutant substitution and the immediate neighbouring residues. A potential transient set of hydrogen bonds between H98 and the polar groups of N123, 123C, 104E and R451, result in the contraction of the secondary structure over a larger extent of the catalytic domain. As a result, the active site surface was reduced in the L98H mutant and the coordination of side chain with the heme prosthetic group differed.

Without revealing free energy reaction pathways of triazole-drug binding, a quantitative measure of how these structural changes affect the binding of drug-to-heme remains beyond this study. However, the results revealed by our simulations at least suggest a qualitative hypothesis of resistance to triazole-drugs within the context of the proteins immediate environment and a direction of work with the models necessary to capture drug pathway mechanisms.

## Acknowledgements

The authors wish to acknowledge the valuable input from Dr. Ana Pabis and Dr. Lynn Kamerlin (Uppsala University). We also wish to acknowledge financial contributions via crowdfunding, which made this work possible: Mrs. Lynda Parnell, Mr James Drinkwater, Mrs. Rachel Drinkwater, Mr. Roland Parkes, Mrs. Jane Parkes, Dr. Julia Halder, Ms. Sarah Vermeulen, Dr. Winnie Wu, Dr. Vince Hall, Mrs. Chandni Rosser, Mr. Ian Wright, Mrs. Jane Shorey, Mrs. Karen Leroy, Mr. Robert Huckle and Mrs. Joanna Huckle.

## Author contributions

A.N. and J.R. conceived and designed the experiments. A.N. performed the experiments. Both authors contributed to the writing of the manuscript.

